# Encoding and Decoding Dynamic Sensory Signals with Recurrent Neural Networks: An Application of Conceptors to Birdsongs

**DOI:** 10.1101/131052

**Authors:** Richard Gast, Patrick Faion, Kai Standvoss, Andrea Suckro, Brian Lewis, Gordon Pipa

## Abstract

In a constantly changing environment the brain has to make sense of dynamic patterns of sensory input. These patterns can refer to stimuli with a complex and hierarchical structure which has to be inferred from the neural activity of sensory areas in the brain. Such areas were found to be locally recurrently structured as well as hierarchically organized within a given sensory domain. While there is a great body of work identifying neural representations of various sensory stimuli at different hierarchical levels, less is known about the nature of these representations. In this work, we propose a model that describes a way to encode and decode sensory stimuli based on the activity patterns of multiple, recurrently connected neural populations with different receptive fields. We demonstrate the ability of our model to learn and recognize complex, dynamic stimuli using birdsongs as exemplary data. These birdsongs can be described by a 2-level hierarchical structure, i.e. as sequences of syllables. Our model matches this hierarchy by learning single syllables on a first level and sequences of these syllables on a top level. Model performance on recognition tasks is investigated for an increasing number of syllables or songs to recognize and compared to state-of-the-art machine learning approaches. Finally, we discuss the implications of our model for the understanding of sensory pattern processing in the brain. We conclude that the employed encoding and decoding mechanisms might capture general computational principles of how the brain extracts relevant information from the activity of recurrently connected neural populations.

## 1 Introduction

How is the brain able to learn about its environment from the sensory input it receives? This fundamental question has been driving neuroscientific research since its beginnings and has been addressed in various domains [19] [32] [42] [18] [28]. Experimental evidence from neuroimaging studies revealed that parts of the brain encode specific sensory stimuli, showing enhanced activity upon presentation of the respective stimulus [21] [34] [13]. Furthermore, similar activation patterns have been observed when subjects had to perform mental tasks involving such stimuli, without the stimuliactually being presented [39] [14]. While this suggests that the brain does form representations of its environment to perform tasks such as recognition, it does not explain how these representations are formed or what information they are based on [37]. Some information on their nature arose from studies showing them to be hierarchically organized in different sensory domains, ranging from simple sensory to more complex and abstract representations [43] [34] [42] [18]. Given such a hierarchical structure, it seems likely that the representations at a given level are based on the activity of neural populations at the preceding level. As it is a known feature of the cortex to be locally recurrently organized [41] [17], the question arises how such representations can be formed based on the activity of recurrently connected neural populations. This question is of particular interest for the case of dynamic sensory input, i.e. signals changing over time. To encode and decode dynamic sensory signals it is necessary to keep some memory of past input, since the input at a given point in time might not be uniquely assignable to a certain signal. While this condition can be satisfied by the membrane potential at the level of single neurons, it can be satisfied by recurrent connections at the level of neural populations. We were particularly interested in the question, whether it is possible to encode and decode multiple dynamic sensory signals with a strongly simplified model of a recurrently connected neural circuit. More specifically, we wanted to investigate whether dynamic signals, as they occur naturally, can be learned by a recurrent neural network (RNN) with very simple neuron models. The crucial task the model has to perform for this purpose, is to extract information about the input to the RNN from its continuously changing activation patterns. This involves observing the state dynamics of the network and detecting patterns in those dynamics that are specific to a certain input signal. Once learned, we wanted to use these representations for recognition of the respective sensory signals. A possible mechanism for the recognition task is proposed by predictive coding theory [12]. It states that internal representations are compared to current sensory input, resulting in a difference signal [3]. This signal in turn is thought to be used to update internal beliefs about which pattern the sensory input might belong to. Therefore, our model needs to be able to both learn representations based on the activity patterns of an RNN and compare already learned representations with the current state of the RNN.

Current advances in the field of reservoir computing provide the tools to build such models. A reservoir is a randomly connected RNN that can be driven with some kind of input pattern and trained to approximate and reproduce that pattern [22]. Each of its neurons has a different, randomly initiated receptive field and a non-linear activation function. In such a network, stimuli are encoded by the state of the whole network, or by a series of states in the case of dynamic input patterns. Unfortunately, reservoirs suffer from so called catastrophic forgetting, which refers to the inability of a single reservoir to learn multiple patterns. However, a recent development by Herbert Jaeger called conceptor can solve that problem [24]. Conceptors rely on the idea that a randomly connected RNN visits only a sub-part of all the possible states it can visit, given an input pattern of limited length. Different inputs should therefore push the network into different parts of its state space, as long as the state space is sufficiently large. A conceptor exploits this behavior by capturing the parts of the state space an RNN visits while being driven with a certain input pattern. It does so by extracting directions of maximum variance from the state development observed in the RNN state space. In other words, conceptors are a dimensionality reduction technique that try to identify the manifolds in the state space of a reservoir in which different signals live. If imposed on the reservoir, the conceptor restrains it to visit only those manifolds, hence acting like an attractor. It is therefore possible to use an RNN of limited size to learn representations of multiple dynamic patterns in the form of conceptors and then use these as input to the same network to reproduce the learned patterns. This makes reservoir-conceptor dynamics a viable option to model representation formation in the brain. Furthermore, it allows for comparing the already learned conceptors to the network dynamics observed for new input to obtain evidence for which pattern the current input might belong to, thus performing recognition.

In the auditory domain the brain has to deal with a one-dimensional, but highly variable and complex dynamic signal. This makes it an optimal candidate to demonstrate pattern learning and recognition within RNNs. In human speech, this complexity is even increased by its deep hierarchical structure, ranging from single phonemes to nested sentences [35] [6]. As capturing this hierarchy would require a similarly deep and complex model, we chose birdsongs as the dynamic pattern to model. Birdsongs have gained increasing popularity for investigating auditory processing in the brain, because of their similar, yet less complex hierarchy as compared to human language, paired with the better understood neural circuits of the auditory bird brain [10] [6]. Many birdsongs show a hierarchical structure combining single syllables to complex songs, thereby resembling human syntax [6]. Such a hierarchical structure has also been found in the bird brain performing song generation [2]. More specifically, high-level neurons encoding syllables fire in a certain sequence that refers to a song and drive neurons on a lower level to initiate the respective motor output [38] [33]. Thus, a hierarchically structured model performing recognition on the level of syllables and songs seems plausible from both a biological and behavioral perspective. Moreover, similar to early language acquisition in humans, song-learning in many bird species has been shown to be error driven [5]. Therefore, song recognition and acquisition processes driven by the similarity between sensory input and stored song representations are readily motivated as underlying computational principles of auditory processing in the bird brain. Birdsongs can be divided in two categories depending on their inherent complexity. Some songs exhibit linear syntax structure, i.e. the song always consists of the same sequence of syllables, whereas other songs show stochastic patterns with the sequence of syllables changing between repetitions. Our study focuses on the linear song syntax found in species like zebra finch and sparrow [36], since it compares better to the deterministic succession of syllables within words in human language.

Recent approaches in modeling birdsongs either focused on mere recognition performance instead of the underlying brain processes [29] [25] [31] or on the brain processes mapping a song to an actual motor output [44] [45]. However, none of these studies tried to capture the inherent hierarchy of the birdsong within a neural network model of the listening bird brain. Here, we propose a birdsong recognition model that recognizes single syllables on a bottom level and sequences of syllables, which we refer to as songs, on a top level. It does so by forming representations of syllables and songs on the respective levels and classifying input according to its similarity to those representations. Both levels are thereby instantiated as reservoirs while syllables and songs are represented by conceptors. By demonstrating that such a model can recognize hierarchical, dynamic signals such as birdsongs, we claim that the recurrent structure at a given level of a certain cortical hierarchy could be sufficient to explain how the brain is able to deal with time-evolving sensory stimuli. Conceptor-learning would thereby describe what kind of information the brain has to extract from its sensory areas to form representations of those stimuli. Namely, it would need to find linear combinations of neurons that describe the directions in the state space of the respective sensory area along which the state of that area varies the strongest under a certain input. Moreover, we propose that conceptors as used for song recognition on the top level of our model are adequate for implementing a predictive coding scheme within an RNN. In summary, we suggest that our model resembles a possible general principle of how the brain can deal with hierarchically structured dynamic patterns based on the activation patterns of recurrently connected sensory areas. The subsequent chapter describes our model in detail, while the third chapter reports the model performance on birdsong data. We believe that a plausible model for recognizing any kind of sensory signal needs to be performing well on basic recognition tasks that the species in question is clearly able to solve. This is why we compared the performance of both levels of our model to state-of-the-art machine learning methods. Finally, we discuss the implications of our model for future research on pattern formation and recognition in the brain as well as for computational models of these processes.

## 2 Model

### 2.1 Syllable Classification Module

Our syllable classification module is based on the model used by Herbert Jaeger for speaker recognition on the Japanese vowel data set [23] with slight adaptations to network parameters and data preprocessing. It consists of a small reservoir in which extracted features of the preprocessed audio data are fed in, as can be seen in the left box of figure 1. The resulting states of the reservoir units are then used to learn conceptors, i.e. representations of each syllable. Those representations can then be compared to the reservoir states emerging from driving the reservoir with test data and thus be used for syllable classification. These two processes are depicted by the green and red arrows of the syllable classification module in figure 1, respectively.

**Fig 1.**
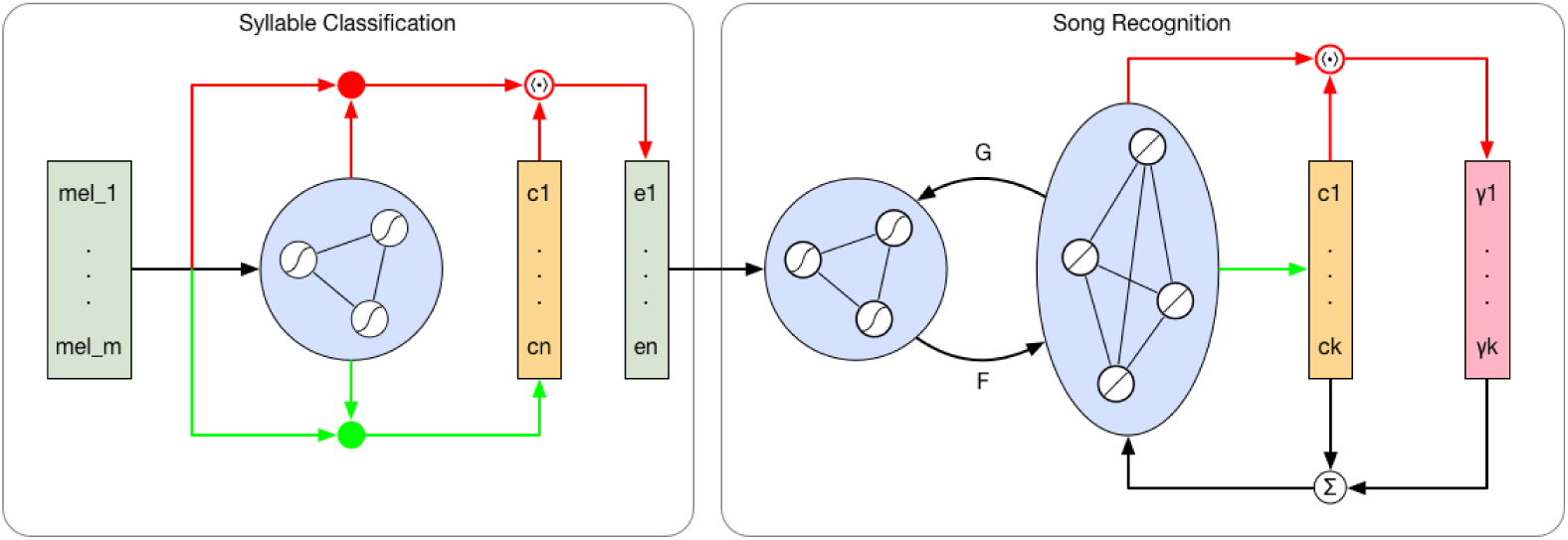
Architecture of the combined model with syllable classification module on the left and song recognition module on the right. Incoming signals are the preprocessed MEL features of the audio signal. Green pathways show information flow for learning, leading to the formation of conceptors *c*. Red pathways show the information flow for classification, leading to evidences *e* or *γ*. Black pathways are for general functionality of the module.

#### 2.1.1 Architecture and Training

The main component of the syllable classification module is a small reservoir of sigmoidal neurons, following the subsequent update equation:

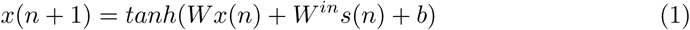

In 1, *x* is the state vector of the reservoir, *W* is the internal connectivity matrix, *W*^*in*^ are the input weights, *s*(*n*) is the input signal and *b* is a bias term. *W* as well as *W*^*in*^ are randomly initialized. On training time, we fed training samples of each syllable into the reservoir and collected its states. We then used those as well as the input of the reservoir to compute a preliminary conceptor *C* for each syllable *j*, employing the following equation:

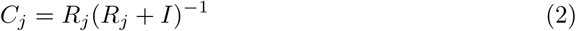

In 2, *R* corresponds to the correlation matrix of the *j*^*th*^ syllable, expressing the correlation of each reservoir unit and input dimension with each other over all training samples, while *I* stands for the identity matrix. On top of that conceptor (which we will from now on refer to as positive conceptor), we also calculated a preliminary negative conceptor for each syllable, resembling all of the state space not occupied by the positive conceptors of all other syllables. This was done using the logical operators for conceptors defined in [23]. With both positive and negative preliminary conceptors at hand, we employed the following equation to receive the final conceptors:

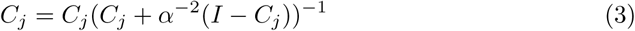

The purpose of 3 is to weave the aperture *α* into the conceptors, which controls how strong the solution is pulled to the identity or to zero. A more detailed explanation of the aperture and how to use it to adapt conceptors can be found in [23]. By the end of this training procedure, each syllable was represented by its own positive conceptor and the logical exclusion of all other syllables.

#### 2.1.2 Classifying Test Data

On testing time, separate test samples of each syllable were fed into the reservoir while collecting its states. Subsequently, we computed the similarity between the collected states *x* for one test sample and each of the conceptors the following way:

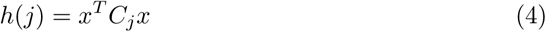

The outcome *h* is a vector containing an evidence value for each conceptor, expressing the similarity between that conceptor and the testing sample that was fed into the reservoir. This was done for the positive and negative conceptors separately. Finally, the resulting evidences were normalized to be restricted to the interval [0,1]. A test sample was then classified as the syllable with the highest combined evidence from positive and negative conceptors.

### 2.2 Song Recognition Module

For song recognition we decided to use an adaptation to the Hierarchical Feature Conceptor (HFC) architecture as proposed by Herbert Jaeger [23]. Similar to the syllable classification module, a conceptor is learned for each song. However, the module does not need to hear the whole song to provide evidences for which song it heard. Instead, it assigns belief values to each learned song and updates them for every new syllable that it receives as input. This update is based on the difference between the network state observed after applying the input and each of the stored conceptors. Another difference to the syllable classification module is the setup of reservoir and conceptors. They follow an architecture which is called Random Feature Conceptor (RFC) and has first been introduced by Herbert Jaeger [23]. This architecture is visualized in the right box of figure 1. In the following, we will explain its sub-parts and dynamics in more detail.

#### 2.2.1 Architecture and Training

The RFC is the main building block of the song recognition module. It is an attempt to store conceptors in a more efficient and biologically plausible way by applying the conceptors on the neurons of a network instead of their connections. This can be done by introducing a second, larger reservoir *z*, whose states evolve in the following way:

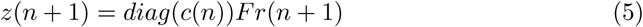

In 5 *diag* refers to the mathematical operator that creates a matrix with the elements of its input vector on the diagonal and zeros on off-diagonal elements. Thus, every row of *F* is weighted by the respective entry in the conceptor *c*(*n*), with *F* being a random matrix mapping the states from the smaller reservoir *r*(*n*) to *z*. The dynamics of *r* are described by:

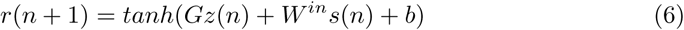

As one can see, 6 is nearly identical to 1 except for the dependency on its own states from the previous time step, which is replaced by a dependency on *z*. *G* is again a random matrix, mapping from *z* to *r*. The dynamics of the RFC as described by 5 and 6 can thus be understood as a loop of two major mechanisms. First, a state space expansion from a smaller reservoir *r* with sigmoidal units to a substantially larger reservoir *z* with linear units performed by *F* and second, a mapping back from *z* to *r* by *G*. In such an architecture, the conceptor *c* can be seen as a single unit or neuron, acting in a multiplicative manner on the units of *z* according to its learned weights to each of them. Thereby, *c* is also time dependent, being updated at each time step according to the following stochastic gradient descent online adaption rule

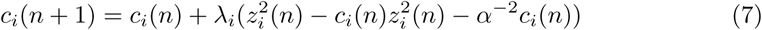

with *λ* being the step size and *α* the aperture. In this case, the aperture determines how strong the conceptor should be able to change in the presence of new information stored in the current state vector *z*. For a more detailed analysis of these dynamics, we refer the interested reader to the chapter on RFCs in [23]. To train conceptors for different songs, we created training patterns of multiple repetitions of each song and fed them into reservoir *r* for a certain training period during which 7 should converge. After this was done for all songs, we trained read-out weights for *r* using Ridge regression over all training patterns [20]. This method chooses the weight for each unit in *r* such that it minimizes the L2 norm of the weights and the sum of the squared residuals between the read-out and the input to *r*. The same method was used to update *G*, changing the internal weights of the mapping from *z* to *r* such that *z* is capable of resembling the input *r* has received during training. This gave us the final RFC which we then used for song recognition.

#### 2.2.2 Classifying Test Data

Song recognition was performed on the timescale of single syllables. More specifically, at every time step a syllable was fed into the network, which returned a belief value for each known pattern, thus performing online song recognition. Thereby, the belief values refer to the weights *γ* assigned to each previously trained conceptor. These weights were used to calculate a weighted sum of all trained conceptors, which was then applied to *z* the same way as the conceptor *c* in 5. They were updated at each point in time according to

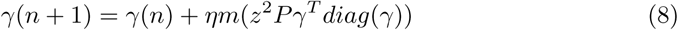

In 8 *η* is the learning rate, *m* is the number of patterns stored in the RFC and *P* is a matrix with a conceptor in each column. Thus, the change of *γ* is a function of the input to *z*, i.e. it is related to the difference between the weighted sum of the stored conceptors and the state of *z* observed for the current input. We used a softmax transformation on *γ* after each update, restricting each weight to [0,1] and ensuring that they sum up to 1. Therefore, each weight can be interpreted as the networks belief in what kind of pattern it is currently driven by. Input was then classified as the song with the highest belief at each time step.

### 2.3 Combined Model

Our final goal was to built a hierarchical model for online birdsong recognition by connecting the syllable classification and the song recognition module, as it is shown in figure 1. In such a model, training had to be done separately within each module, as the conceptors used for syllable representations cannot be learned from the error signal on the top level as of yet. We started out by creating a set of songs (i.e. a set of syllable sequences) from a pre-defined set of syllables. We then trained our syllable classifier on each syllable and ran the songs through it, receiving sequences of syllable evidences *h*. Those syllable evidences are a noisy representation of the original songs and can be interpreted as beliefs of lower level sensory areas in the type of syllable heard. Those noisy representations were then used to train the RFC on each song in the above described way. On testing time, we were again running our syllable test samples through the syllable classifier. The resulting syllable evidences were used afterwards to drive the song recognition module. This gave us a weight *γ* for each song at each time step, indicating the evidence for which song is sung at a given moment. The full data flow of the combined model can be seen in figure 2.

**Fig. 2.**
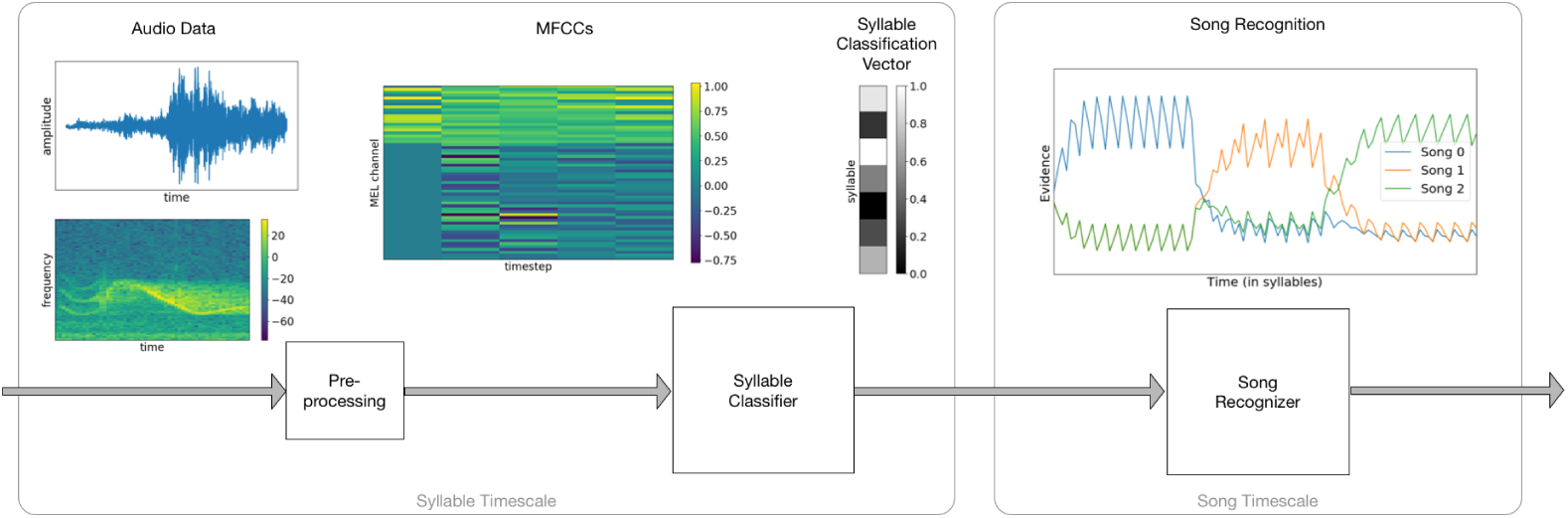
Dataflow visualization for the combined model. The left side is processed on syllable timescale, i.e. in seconds, the right is processed on song timescale, i.e. in syllable indices. Raw audio data from syllables is preprocessed into enriched MEL features and classified by the syllable classifier. The classification vectors are fed over time into the song recognizer, which produces an evidence value for each of the learned songs.

## 3 Model Evaluation

### 3.1 Dataset

Our model was tested on a combination of bird recordings and synthetic data. All of our syllable data was extracted from the online database BirdDB [1]. This database contains recordings from Cassin’s Vireo (*vireocassini*) among other bird species, including syllable annotation for a total repertoire of 65 different syllables. We cut the recordings according to the syllable annotations, creating sets of samples for each syllable. Such a recording is visualized for an exemplary syllable in figure 2. These syllable samples were down-sampled to 20 kHz. Afterwards, we extracted 20 mel frequency cepstral coefficients (MFCCs) as these have not only proven to be very useful for human speech recognition, but also for birdsong classification [30]. The extracted MFCCs were then normalized to a total of 4 time steps and each coefficient was bound to the interval [0,1]. In a final step, we added the first and second derivative of each coefficient. Thus, every syllable was represented by a vector of 60 spectral features at each time step as it is depicted for a single syllable in figure 2. The input *p* to the syllable classification module as in 1 were those spectral feature vectors. As the Cassin’s Vireo does not have a linear song syntax, we could not use these data for song recognition, but only for syllable classification. Therefore, we created a total of 20 synthetic songs from the set of 65 syllables. Each song consisted of 3-5 syllables and was of Markov order 1 or 2, as it is typical for birdsongs [9]. In this case, the Markov order refers to the number of previous syllables necessary to predict the next syllable within a particular song. We created a number of songs for each combination of song length and Markov order using different sets of syllables for the songs of each combination. A song was represented by *n* vectors, whereas *n* refers to the number of syllables the song consists of. Such a vector was of length *m* with *m* being the number of unique syllables present in the set of songs used for the current simulation. Each entry in that vector referred to a single syllable and represented the evidence for that syllable being the input at that particular time-step. A noise free syllable representation would thus be a vector with all entries being 0 except the one of the syllable currently being input, which would be a 1. These syllable vectors were then used as input *s* to the song recognition module as in 6.

### 3.2 Results

The following paragraphs describe how well our model performs on the tasks it was trained on. To evaluate the performance of a model on a task, some kind of comparison or benchmark is necessary. When modeling a sensory recognition task, ideally one would like to compare against behavioral data from actual living organisms performing the same task. Unfortunately, to our knowledge there is no data on how well certain birds are able to recognize bird songs or syllables under different conditions like increased sensory noise. Therefore, we decided to compare our model against other methods for bird song or syllable classification. We did not compare against performance values reported in the birdsong recognition literature, since most of these values refer to species recognition or were performed on substantially different data sets. Instead, we decided to implement alternative methods for song and syllable recognition ourself. We chose to use deep neural network (DNN) architectures as comparison to our model, since they have proven to be the best performing supervised methods on time series data like human speech [40] [7]. Due to the available training data being limited, we used rather simple DNN architectures, which showed convergence behavior in their performance on the training data sets. We will first compare the classification performance of our bottom level syllable classifier with a multi-layer perceptron (MLP) on the syllable classification task and then compare the performance of our top level song recognition module with a gated recurrent unit (GRU) network on an online song recognition task. Finally, we will evaluate the behavior and performance of our combined model during song recognition.

#### 3.2.1 Syllable Classification

We tested the performance of our syllable classification module on an increasing number of syllables to distinguish between. Furthermore, we also tested a shallow MLP on the same task, serving as a comparison to our conceptor-based syllable classification. It employed a single hidden layer of hyperbolic tangent units whose size matched the number of input features. With 20 MFCCs plus their first and second derivatives evaluated at 4 time steps, this amounted to a total of 240 units. The output layer used a softmax activation function and its size always equaled the number of unique syllables the MLP was trained on. During training, the cross-entropy between MLP output and target was minimized using an Adam optimizer with an initial learning rate of 0.001 [27]. Initial weights were sampled from a normal distribution with zero mean and standard deviation of 0.1. We trained the MLP on all training samples for 3 epochs using mini-batches of size 10. The reservoir of the conceptor-based classifier was initialized with 10 neurons. The apertures for positive and negative conceptors were *α*_*p*_ = 25 and *α*_*n*_ = 27, respectively. Both models were trained and tested 10 times for each number of syllables within the range from 5 to 55. Each run used a different set of randomly drawn syllables and random training-test data splits with 50 training samples and 10 test samples. Each test sample was classified as the syllable with the highest evidence from the respective output units. We then calculated the overall classification performance as the mean of correct classifications over all test samples. The resulting mean classification performances of conceptor-based and MLP classifier are visualized in figure 3 A. As can be seen, the performances of the classifiers are very similar for smaller syllable numbers and drop off with increasing syllable numbers. Thereby, performance drops slightly faster for the conceptor-based classifier. However, for the amount of songs we test our full model on, the number of used syllables never reaches a regime in which the MLP clearly outperformes the conceptor-based classifier. In part B of figure 3 the evidences for a single test run with 30 syllables are depicted. It is worth noticing how the evidences of the negative conceptors clearly reduce the number of miss-classifications. This effect was observed for 73,6% of our test runs and demonstrates the potential of logical operations for cognitive tasks such as inferring the nature of sensory input.

**Fig. 3.**
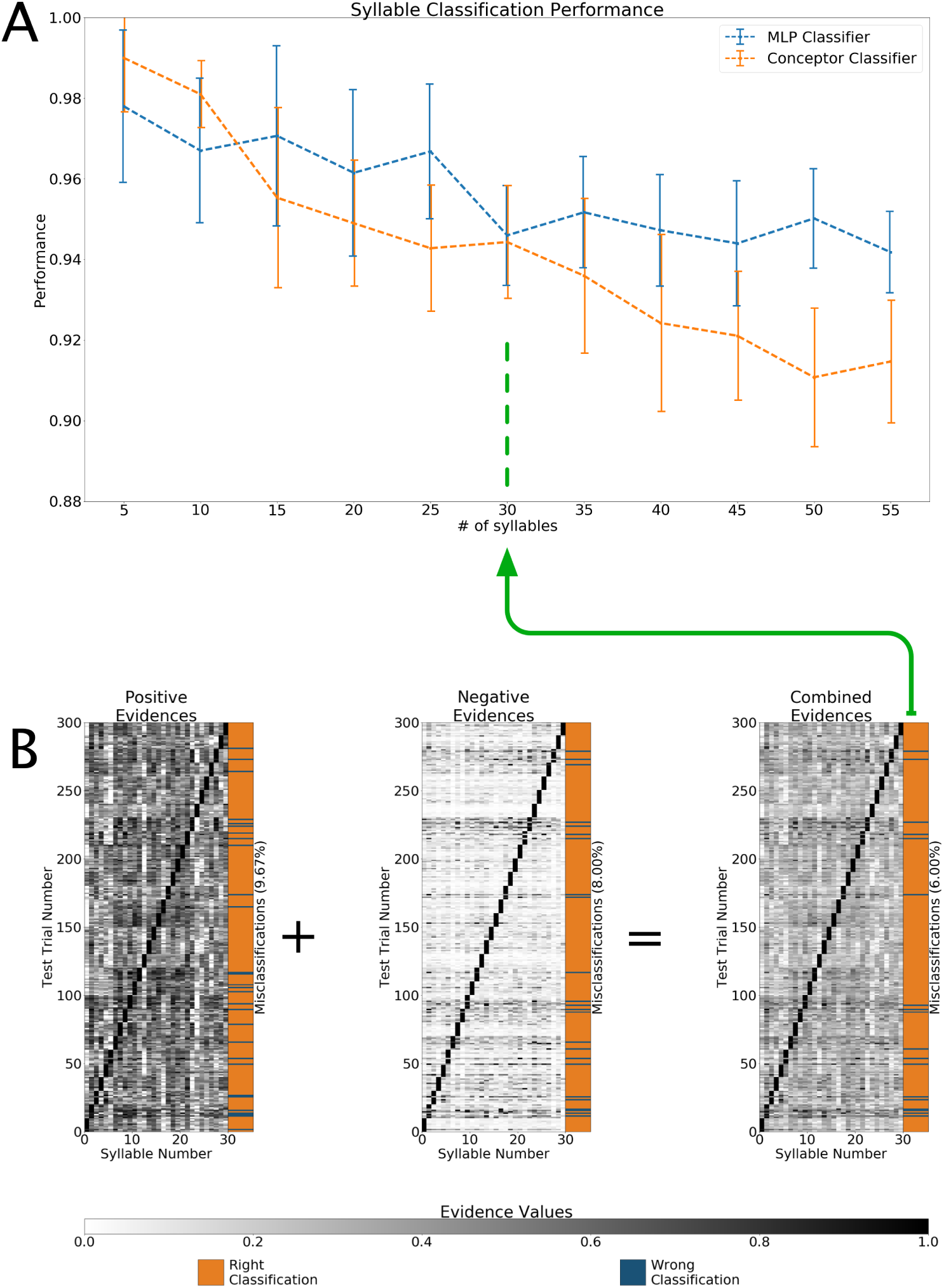
A: Mean classification performance of the conceptor-based syllable classifier and a multi-layer perceptron on different numbers of syllables. Error bars indicate the standard deviation over 10 trials. B: Evidences from positive, negative and combined conceptors for an exemplary test run with 30 different syllables.

#### 3.2.2 Song Recognition

The performance of our song recognition module was also investigated for an increasing number of songs. However, within the full model the module will have to learn and recognize songs based on the input of the syllable classifier. The noise level of this input will depend on the number of used syllables, as we have learned from the first experiment. Therefore, we additionally tested the robustness against noise of our song recognition module by adding white noise of variable strength to each of the syllable vectors fed to it on testing time. We tested the song recognition performance for up to 7 different songs and a signal-to-noise ratio (SNR) ranging between 4:1 and 1:8. For each combination of song number and SNR we compared the performance of our conceptor-based classifier against a GRU network. The latter consisted of 3 hidden layers, each of which contained 100 units with rectified linear activation. All 3 hidden layers were regularized by dropout of 40%. As input the syllables comprising the songs were fed to an embedding layer, converting the scalar index representation into a vector of length 16. The output was calculated using a softmax layer with the number of units equaling the number of different songs. The categorical-crossentropy between output and targets was minimized using an Adam optimizer with initial learning rate of 0.001. The network was trained for 10 epochs with a mini-batch size of 64. Hyper-parameters were optimized using Tree of Parzen Estimators implemented in the hyperopt package [4]. The conceptor-based classifier was initialized with a smaller reservoir of 400 neurons and a larger reservoir of 2000 neurons. We chose learning rates of *λ* = 0.5 and *η* = 0.005 and an aperture of *α* = 3. Both classifiers were trained and tested on randomly selected sub-sets of the 20 synthetic songs described above. Thereby, the training set consisted of a total of 600 song samples while the test set consisted of 100 song samples. On test time, we collected the classifiers’ outputs for each input vector and used a winner-takes-it-all transform on those to arrive at a song classification. Performance was then measured as the fraction of correct classifications over the entire test data set. Each combination of song number and signal-to-noise ratio was tested 10 times with different randomly selected songs. The mean results of those simulations are visualized in figure 4. For our song recognition module we can observe the effects of song number and SNR on recognition performance to be roughly additive, as indicated by the diagonal structure of the performance matrix. This is very different from the GRU classifier, which can deal well with increasing numbers of songs, but is more susceptible to noise. Looking at the difference between the two performance matrices, it is clearly visible that the conceptor-based classifier is superior at low SNRs. It is important to notice that it was trained on noise-free training data, while we added noise to the training data of the GRU to improve its performance. This training data noise was in the range of the noise used for the testing data. We do recognize that increasing the amount of training data could have improved the performance of the GRU classifier. However, since both methods are supervised and we already allowed the GRU to see noisy training data, we decided to keep the amount of training data equal for both methods. While these results demonstrate that the combined model should be robust to the noise of the syllable classifier on test time, it remains unclear how well the song recognition module can deal with noisy training data.

**Fig. 4.**
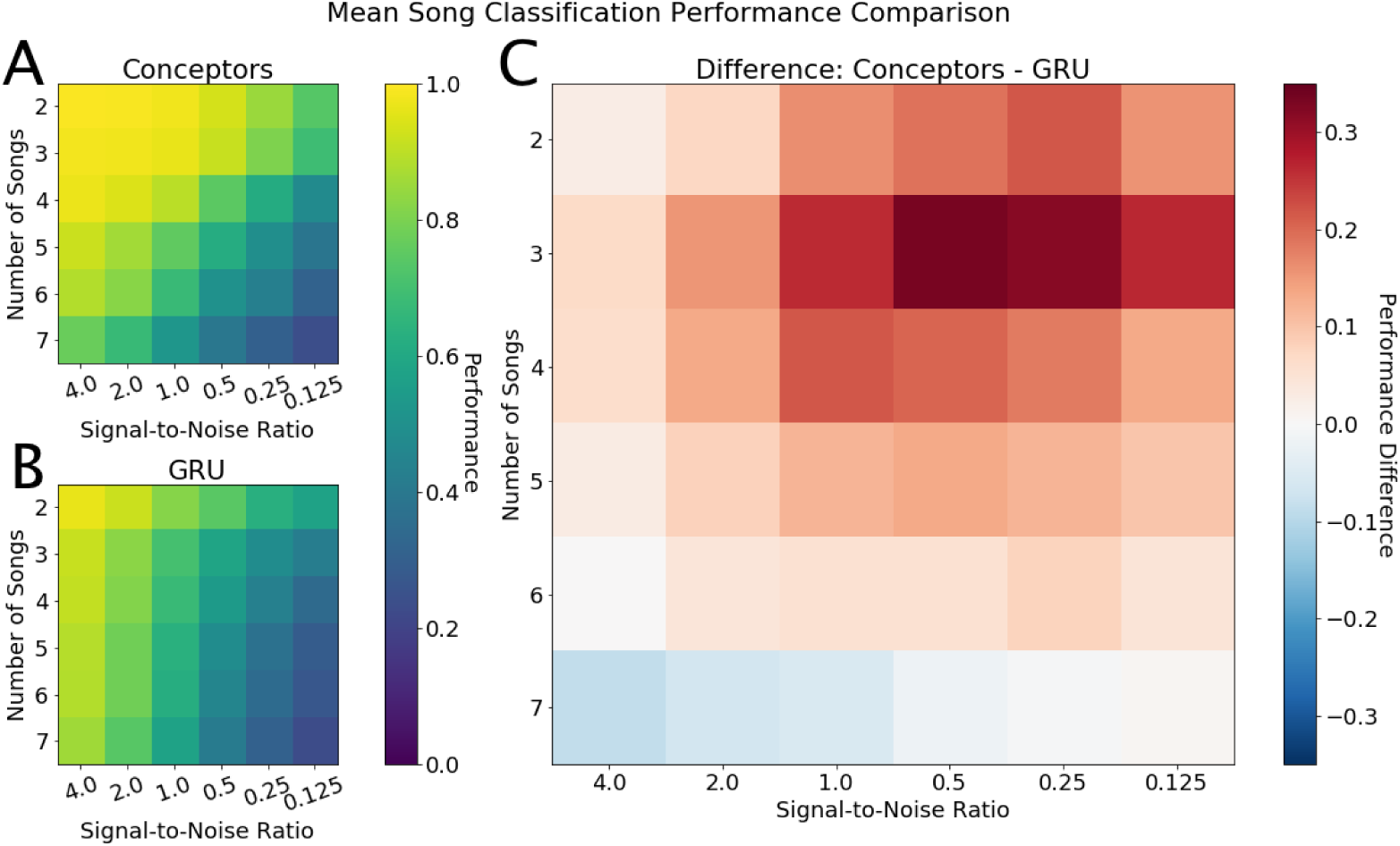
Left: Mean classification performance of conceptor-based classifier and GRU classifier on different numbers of songs and signal-to-noise ratios. Right: Differences in classification performance between both classifiers

#### 3.2.3 Combined Model

In a final step, we investigated whether the song recognition module is able to learn and recognize songs based purely on the syllable evidences provided by the syllable classification module. Again, we tested the performance for up to 7 songs drawn randomly from the 20 synthetic songs described above. The syllable classifier was trained on all syllables used for the songs of a respective run. Afterwards, we created a training and a test data set consisting of multiple repetitions of the randomly drawn songs. The training data consisted of around 400 repetitions of each song, while the test data consisted of a total of 100 song repetitions. We then drew a random wave sample for each syllable in training and test data, using different sets of samples for the both. Subsequently, we ran these data through the syllable classifier and stored the resulting syllable evidences as final training and test data. The song recognition module was then trained on the respective songs. In a final step, we drove the fully trained model with the test data and collected the yielded song evidences. This procedure was repeated 10 times for different random splits of training and test samples and different randomly chosen songs. Model parameters were set to the same values as reported in the results sections of the single modules. The recognition performance for 2 to 7 different songs can be observed in part A of figure 5. As evidenced by the position of the performance values within the performance range of the song recognition module, the combined model performs as well as the mere song recognition module on high SNR test data. Thus, the song recognition module proved to be robust to the noisy data provided by the syllable classifier for both song learning and song recognition. In part B of figure 5 the dynamics of the song recognition process are depicted for an exemplary test run of multiple repetitions of 3 different songs. During the time course of a single song repetition the belief value for the respective song increases while the belief values of the other songs decrease. At the end of a song belief values tend to go back to chance level given no new input, meaning that pauses between song repetitions function as reset if sufficiently long. Within part B of figure 5 there are no pauses, i.e. the end of a song repetition is followed by another song immediately. In cases where the following song is the same as the previous song, the belief values stay within a similar range, while they adapt very quickly (within a single song repetition) given a new song. Thus, miss-classifications usually only happen at the very beginning of the initial repetition of a new song.

## 4 Discussion

The main goal of our work was to demonstrate how a recurrent neural circuit can encode and decode complex dynamic patterns such as birdsongs. Thereby, we employed a recently developed technique that allows the capture and control of the dynamics of randomly connected RNNs [23]. Using this mechanism, we were able to show how birdsongs can be learned and recognized within a 2-level hierarchical model of RNNs. While the first level learned and recognized single syllables based on their spectral features, the second level learned and recognized songs as sequences of syllables. Both processes, learning and recognition, relied on extracting information from the activity pattern of an RNN driven by dynamic input. We were able to show that on a limited number of distinguishable patterns the recognition performance of our model compares to state-of-the-art machine learning methods and that song recognition is remarkably robust to noise, a desirable property when dealing with recognition in natural environments. Our model was motivated by two basic organizational properties of the brain. First, its recurrent structure on the level of local neuronal circuits [8] [41], and second, the hierarchical organization of the brain and the representations stored therein [12]. If the description of sensory areas in the brain as recurrently connected recurrent neural circuits is sufficient to capture the crucial information transmitted by those areas, our model suggests general computational principles for encoding and decoding dynamic sensory signals in the brain.

It is important to emphasize that our model merely describes what kind of computations the brain has to perform given a certain architecture and how they can be implemented by means of reservoir computing. It does not try to answer how those computations are performed by the brain. The smallest units of our network are sigmoidal neurons, which can resemble the mean activity of a population of real neurons at best. As this is a hugely oversimplified model of neuronal computations, our model cannot resemble the complex processes at work in the brain during representation formation. Despite these limitations, it can still describe what influence a certain sensory input has on a recurrently connected neuronal population and how the resulting change in the state of that population can be used to encode and recognize patterns of different complexity. Thus, while we do not claim that sensory representations in the brain are formed in any way like the conceptors used in our model, we nevertheless propose that they could contain the same information as captured by the conceptors. Specifically, we highlight the importance of extracting information about the state development of a neural circuit and about which directions of the state space are important and which ones are unimportant in describing the state development under a certain input. In other words, we believe dimensionality reduction to be an essential tool for extracting information from the activation patterns of large neuronal populations in hierarchically organized neural circuits. Conceptors are a special form of dimensionality reduction and we demonstrated its usefulness for dynamic sensory pattern recognition within this paper. Furthermore, our model shows how such pattern recognition processes could be embedded in a hierarchical structure. This hierarchy is not only present in the topography of the model, but also in its timescales. While the bottom level reservoir develops on the timescale of the feature representation we chose for our syllables, the top level evolves on the timescale of full syllables. As each syllable is represented by spectral features at 4 different time steps, the top level reservoirs evolve approximately 4 times slower than the bottom level reservoir. This behavior is in line with experimental evidence on timescales in the brain [26].

**Fig. 5.**
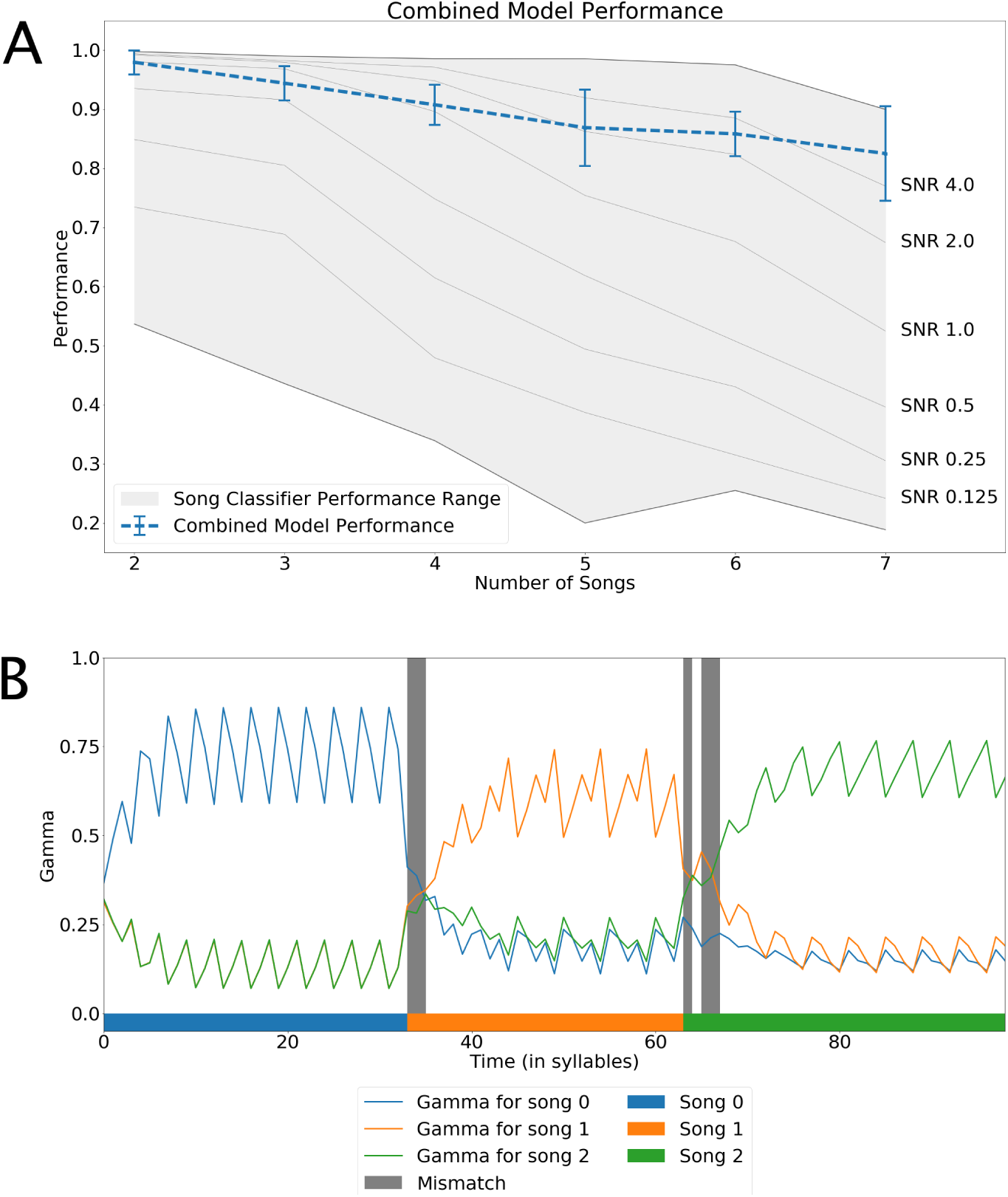
A: Recognition performance of the combined model for different song numbers in the context of the song recognition performance range. B: Song evidences (*γ* values) for an example run of the song classifier for multiple repetitions of three different songs.

Given that descriptive level, we showed that to learn a stimulus based on the activation pattern of a neuronal population, the brain has to find out which linear combinations of neurons encode information about the stimuli and store that knowledge somehow. In our model, this process is represented by creating the conceptors which amplify neurons whose activity varies a lot over the course of an input pattern while suppressing the ones that do not. On top of that we showed how recognition of new stimuli could be performed given such stored knowledge about stimulus patterns. This has been done differently in song and syllable classifier, however. For song classification, we implemented a predictive coding scheme employing random feature conceptors. Using RFCs as song representations comes with two desirable properties. The first one is related to the weights *γ* assigned to each conceptor. These weights formed a probability distribution, expressing the networks current belief in what song it was driven by. At a given time step, this belief was a function of the belief from the previous time step as well as the new evidence gathered from the changes in the network states caused by new input. This dependency is in line with Bayesian inference steps performed within a predictive coding framework [28]. Since each conceptor in our top-level module was a unit with weights to every neuron in the reservoir *z*, they could also be interpreted as single neurons with synaptic connections. From such a perspective the weights *γ* would represent the activation of the neurons, meaning that each neuron’s activity would be determined by the similarity of the song it codes for to the current input to the reservoir *r*. Such activation could then be detected by a hierarchically even higher level and used for decision processes and the like. This could be a possible way of how probability distributions are represented and used in the brain. Furthermore, this offers a direct way to implement top-down influences by acting on the weights of the conceptors. The second desirable property is that the trained conceptors can serve as generative models. More specifically, applying a conceptor encoding a certain song to the RNN it was trained on will force the RNN to generate that song without any further input needed. Therefore, one cannot only use the learned representations for recognition, as demonstrated in this paper, but also for other cognitive tasks requiring generation processes. One drawback to using RFCs is the limited stability of the training process. At its current development state, the RFC can sometimes fail to converge to a correct solution for a certain pattern. More specifically, it can fail to capture the subspace of the RNN state space which the RNN visited for a certain input pattern. This is not a problem for the presented recognition task, as once a correct conceptor is learned for every pattern, the network performance is stable. Unfortunately, the probability of at least one pattern not being learned correctly increases with the number of overall patterns to learn. Therefore, given limited training data and computational capacities, testing recognition performance for more than 7 songs was unfeasible for us. To resolve that issue in a satisfactory manner, more work has to be invested in stabilizing the RFC architecture. One possible direction of research could be, to extend conceptors to other dimensionality reduction mechanisms apart from the one currently employed in conceptor learning.

Due to that limitation of the RFC we used a more simplified architecture for syllable classification, which allowed us to use a bigger set of syllables to construct our songs from. While the conceptors within this simplified architecture cannot serve as generative models, they still allow for using the logical operations *AND OR* and *NOT* on them. Thus, we were able to calculate negative conceptors for each syllable, representing the logical exclusions of all other syllable conceptors. Combining the evidences of the network being driven by a certain syllable with the evidence of the network not being driven by any of the other syllables boosted our classification performance significantly. Integrating these logical operations into an online recognition process such as the song recognition on the top level would in theory be possible as well. However, one would have to integrate both positive and negative conceptor into a single one in order to use the update scheme for the belief values in the way it is laid out in this paper. Importantly, the simplified architecture of the syllable classifier sufficed to demonstrate that an abstract spectral representation of highly variable sound patterns such as bird calls is sufficient to learn and recognize those syllables. This is in line with experimental evidence suggesting that mental representations are typically under-specified and abstract, making them more robust to noise [11].

In summary, we put forth a hierarchical model of RNNs which is able to recognize non-stochastic birdsongs online. However, the mechanisms used in the model are in principle not specific to that one task or domain. Every dynamic pattern of similar length and dimensionality could be learned and recognized by such an architecture. Thereby, the complexity of the hierarchy of the input could be met by a suitable amount of hierarchical levels in the model. Our model suggests a general principle of how the activity pattern of a population of neurons with different receptive fields could be used to encode and decode information about and from the environment. Moreover, it shows how these principles could be implemented in a hierarchically organized neural circuit performing recognition tasks, though they could even be used in a similar fashion to perform pattern generation tasks.

